# Single-cell analyses reveal that monocyte gene expression profiles influence HIV-1 reservoir size in acutely treated cohorts

**DOI:** 10.1101/2024.11.12.623270

**Authors:** Philip K. Ehrenberg, Aviva Geretz, Meta Volcic, Taisuke Izumi, Lauren Yum, Adam Waickman, Shida Shangguan, Dominic Paquin-Proulx, Matthew Creegan, Meera Bose, Kawthar Machmach, Aidan McGraw, Akshara Narahari, Jeffrey R. Currier, Carlo Sacdalan, Nittaya Phanuphak, Richard Apps, Michael Corley, Lishomwa C. Ndhlovu, Bonnie Slike, Shelly J. Krebs, Jintanat Anonworanich, Sodsai Tovanabutra, Merlin L. Robb, Michael A. Eller, Gregory M. Laird, Joshua Cyktor, Eric S. Daar, Trevor A. Crowell, John W. Mellors, Sandhya Vasan, Nelson L. Michael, Frank Kirchhoff, Rasmi Thomas

## Abstract

Elimination of latent HIV-1 is a major goal of AIDS research but the host factors determining the size of these reservoirs are poorly understood. Here, we investigated whether differences in host gene expression modulate the size of the HIV-1 reservoir during suppressive ART. Peripheral blood mononuclear cells (PBMC) from fourteen individuals initiating ART during acute infection who demonstrated effective viral suppression but varying magnitude of total HIV-1 DNA were characterized by single-cell RNA sequencing (scRNA-seq). Differentially expressed genes and enriched pathways demonstrated increased monocyte activity in participants with undetectable HIV-1 reservoirs. *IL1B* expression in CD14+ monocytes showed the greatest fold difference. The inverse association of *IL1B* with reservoir size was validated in an independent cohort comprised of 38 participants with different genetic backgrounds and HIV-1 subtype infections, and further confirmed with intact proviral DNA assay (IPDA^®^) measurements of intact HIV-1 proviruses in a subset of the samples. Modeling interactions with cell population frequencies showed that monocyte *IL1B* expression associated inversely with reservoir size in the context of higher frequencies of central memory CD4+ T cells, implicating an indirect effect of *IL1B* via the cell type well established to be a reservoir for persistent HIV-1. Signatures consisting of co-expressed genes including *IL1B* were highly enriched in the “TNFα signaling via NF-κB” geneset. Functional analyses in cell culture revealed that IL1B activates NF-κB, thereby promoting productive HIV-1 infection while simultaneously suppressing viral spread, suggesting a natural latency reversing activity to deplete the reservoir in ART treated individuals. Altogether, unbiased high throughput scRNA-seq analyses revealed that monocyte *IL1B* variation could decrease HIV-1 proviral reservoirs in individuals initiating ART during acute infection.

## INTRODUCTION

HIV-1 infection is effectively treated with antiretroviral therapy (ART). However, the persistence of stably integrated and replication-competent proviruses in the latent cell reservoir prevents a cure ^1^. ART suppresses plasma viremia below the limit of detection but viral replication rebounds within weeks of analytic treatment interruption (ATI) in the majority of individuals ^2^. This is attributed to a small pool of latently infected cells harboring HIV proviruses, which can be reactivated to produce infectious viruses that cause viral rebound in the absence of ART ^3^. Several studies have implicated resting memory CD4+ T cells and distinct memory CD4+ T cell compartments as the primary latent reservoirs in people living with HIV (PLWH) on ART ^4–7^. In long-term non-progressors (LTNP) and elite controllers (EC), viremia is controlled in the absence of treatment due to the presence of protective HLA alleles including B*57; however, there exists a third considerably smaller group of post-treatment controllers (PTC) without these protective alleles, suggesting a distinct mechanism of viral control ^8,9^. Although a recent clinical trial identified one controller who had HLA-B*57 ^10^ in the placebo study arm after ATI, perhaps the most consistent correlate of post-treatment control is the presence of lower cell-associated HIV proviral DNA levels as a surrogate of CD4+ T cell reservoir size^9^.

Viral rebound is observed after treatment interruption in almost all people, whether ART is initiated at the early acute or later chronic stage of HIV infection ^11,12^. However, there is increasing evidence that the pool of latently infected cells, which persist despite treatment, varies in size between individuals. Variation in reservoir size as determined by HIV DNA quantification in CD4+ T cells has also been observed in virally suppressed patients who initiated ART during acute HIV infection ^13,14^. This inter-host variation in CD4+ T cell associated reservoir size is observed at various stages of acute infection and even after 24 weeks of ART ^13,15^. Identifying host cellular factors that mark and influence the HIV reservoir size could help in understanding the mechanisms associated with HIV persistence and may reveal targets for achieving a functional cure. The majority of previous findings linking the host transcriptome to latency have been limited to cell lines or models of infection, and ex vivo experiments with primary cells ^16–18^. Recent studies in humans have focused on assessing CD4+ T cells and HIV persistence in the context of characterizing antigenicity, clonal expansion, and the whole transcriptomes of single cells harboring virus ^19–22^. Here, we determined cellular immune profiles of the host that associates with cell-associated HIV DNA levels, an established marker of viral persistence using extreme phenotypes of reservoir size ^23^. These in vivo quantitative phenotypes of multiple donors enable unbiased approaches to interrogate all cell populations without ex vivo stimulation. A unique cohort of PLWH that initiated ART treatment during Fiebig stage III of acute infection was selected in order to minimize the effect of time-to-treatment as a confounder of reservoir size ^24^. This was combined with the use of analytical approaches that are unbiased and high throughput to avoid specifically targeting the known latently infected T cell reservoir, and to enable broad screening for host variation most prominently associated with reservoir size.

Given the sustained size variation in cellular reservoirs during acute HIV infection (AHI) and post-ART initiation, we hypothesized that specific host genes might contribute to these differences between individuals. We have previously shown that a bulk RNA-seq approach applied to multiple specific sorted lymphocyte populations allowed us to identify protective gene signatures in response to HIV vaccination ^25^. Here, we used a more sensitive next-generation sequencing (NGS) approach to identify differences in host transcriptomes from PLWH shown to harbor varying HIV DNA levels ^24^. Additionally, we recently showed that transcriptomics studies conducted with AHI samples can be confounded by the presence of viremia ^15,26^. Here, we achieved broader scope and resolution using an unbiased droplet based single-cell RNA-seq (scRNA-seq) platform with peripheral blood mononuclear cell (PBMC) samples from virally suppressed PLWH. This enabled the examination of gene expression in all cell types in peripheral blood, to test expression of every gene in the transcriptome from individual cells for associations with the size of the total cell-associated HIV reservoir in donors on ART. Our unbiased single-cell analyses identified monocyte gene expression profiles as having the strongest association with HIV reservoir size in two independent AHI cohorts, with significant expression of higher *IL1B* with a smaller reservoir in both studies. Moreover we were able to confirm these findings with IPDA^®^ measurements of intact HIV-1 proviruses. Functional in vitro data support an effect via IL1B-mediated activation of the transcription factor family NF-κB, which both stimulates HIV transcription and induces antiviral gene expression ^27^.

## RESULTS

### Participant selection and study design

We screened 163 Thai participants enrolled in the RV254 AHI cohort with varying cell-associated HIV-1 DNA reservoir sizes to map the immune-microenvironment of PLWH. We further focused on performing integrated transcriptome-wide scRNA-seq, high parameter multidimensional flow cytometry, T-cell receptor (TCR) and B-cell receptor (BCR) sequence data from PBMCs of 14 selected participants, who had been on ART for 48 weeks. All individuals had initiated ART following HIV diagnosis during Fiebig stage III of AHI and were virally suppressed (<50 copies/ml) (Extended table 1). We categorized the 14 participants with the most extreme reservoir size differences into “undetectable” (n=8) versus “detectable” (n=6) reservoir groups from a total of 28 participants at Fiebig stage III (Fig. 1A-B). These phenotypes were based on both total cell associated and integrated HIV DNA levels in PBMC as measured by quantitative PCR ^14,28^. Further, HIV DNA decay from week 0 at AHI showed that the phenotype categorizations were distinct (Fig. 1C). All 14 participants shared similar demographics and were Thai males infected with viral subtype CRF01_AE, as described previously ^22^.

**Figure 1.**
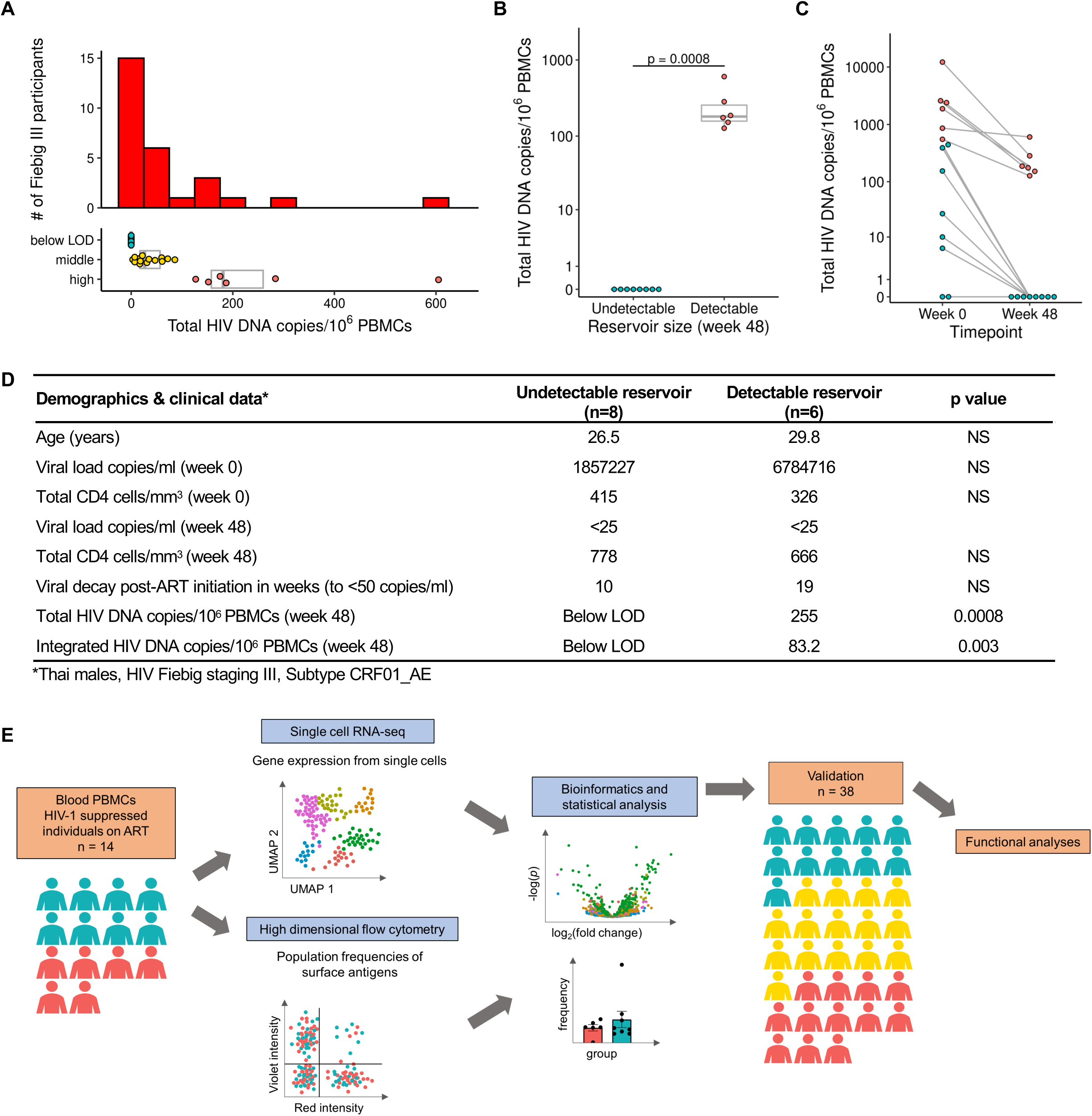
Characteristics of study participants and experimental design. A) Distribution of total HIV DNA in Fiebig stage III participants in RV254 at week 48 after ART initiation and their categorization into three groups based on reservoir size. B) Selected participants from Fiebig stage III with extreme reservoir size phenotypes (undetectable = below LOD and detectable = high) of cell-associated total HIV DNA in the RV254 Thai discovery cohort (n=14). Significance was determined by the Mann-Whitney U test. C) Total HIV DNA decay between weeks 0 (AHI) and 48 (after ART initiation). D) Phenotypes of participants comprising the detectable versus undetectable reservoir size categories. Mean values are shown for each group, NS: not significant. E) Single-cell RNA-seq and multiparameter flow cytometry were performed on all 14 participants. Additional validation by scRNA-seq was performed in an independent AHI cohort from the USA (A5354) (n=38).

Other than reservoir size there were no significant differences between the two groups (Figs. 1B, D). The workflow including scRNA-seq, repertoire sequencing, and flow cytometry performed on samples from all 14 donors is illustrated in Fig. 1E. Furthermore, PBMC from an additional 38 male participants with viral subtype B infections and African and European ancestry from the USA (ACTG A5354) were assessed 48 weeks after ART initiation for validation of cell subset-specific differential gene expression patterns with reservoir size (Fig. 1E, Extended table 1).

### CD14+ monocytes have the most differentially expressed genes associated with reservoir size

PBMC from the 14 Thai male participants collected 48 weeks after ART initiation were assessed by scRNA-seq on the 10x Genomics platform using 5’ gene expression profiling. A total of 62,925 single cells passed quality filter and 19,581 genes were detected across all cell types from all donors. Cell clustering based on gene expression of lineage markers revealed 24 discrete populations (Figs. 2A, S1). All major canonical immune cell populations in PBMC could be detected through gene expression, including cells from the innate, humoral and cellular arms of the immune system (Supplementary table S1). There were no significant differences in uniform manifold approximation and projection (UMAP) distributions or cell subset frequencies when comparing detectable versus undetectable reservoir groups (Figs. S2A-B). Furthermore, no apparent differences in T cell receptor (TCR) or B cell receptor (BCR) clonal diversity or in BCR isotype distribution were observed between detectable and undetectable reservoir groups across all conventional T and B cell subsets captured in this analysis (Fig. S3A-C). We performed differential expression analyses to identify genes whose expression showed quantitative differences between people with undetectable or detectable amounts of HIV DNA 48 weeks after ART initiation in all 14 participants. These analyses identified significant differences in gene expression between the two groups in 20 cell subsets. There were 224 unique significantly differentially expressed genes (DEG) which were independent of the size of the immune cell subsets. The cell types with the highest number of DEGs were CD14+ monocytes (n=78), CD8+ T_CM_ cells (n=51), CD8+ T_EM_ cells (n=46) and CD16+ monocytes (n=38) (Fig. 2B, Supplementary table S2). Analyses of the 224 DEG identified the top 20 significant pathways and processes collectively enriched across the different cell subsets (Supplementary table S3). The top 3 significant gene ontology terms were lymphocyte activation, regulation of cytokine production, and cytokine-mediated signaling, with most of the pathway enrichments resulting from the DEG in monocytes. The only pathways that were significantly enriched in the detectable reservoir group were lymphocyte activation and immune response-activating signal transduction in CD4+ naïve T cells (Supplementary table S4). DEG with the greatest significance in different cell subsets are highlighted in Fig. 2C. The DEGs that were most significant, with an average log fold change of >1, were thrombospondin-1 (*THBS1*) and interleukin-1 beta (*IL1B*) in CD14+ monocytes (Fig. 2C). The median expression of these two genes in CD14+ monocytes was significantly higher in the undetectable compared to the detectable reservoir group when assessed using a single-cell approach (P_adjusted_<5e-324 and P_adjusted_=8.4e-197 for *THBS1* and *IL1B*, respectively) or by participant-specific average gene expression analyses (P=0.02 and P=0.001 for *THBS1* and *IL1B*, respectively) (Fig. 2D-E). These genes were consistently expressed at higher levels for individuals with undetectable reservoirs, whether measurements were determined by total HIV DNA in PBMC or only in CD4+ T cells (Figs. S4A-B). The associations remained significant even when the outcome was HIV reservoir decay from week 0 (AHI) to week 48 (on ART) (Coefficient = 4.99e-04 P_adjusted_ <5e-324 and Coefficient = 1.47e-04, P_adjusted_=3.97e-104 for *THBS1* and *IL1B*, respectively (Fig. S4C-F). Thus, from an unbiased screen of all peripheral blood cell populations, we observed the strongest correlations with reservoir size not for CD4+ T cell subsets, but for monocytes, which showed enrichment for activation pathways, and particularly increased expression of *THBS1* and *IL1B*, in the undetectable reservoir group.

**Figure 2.**
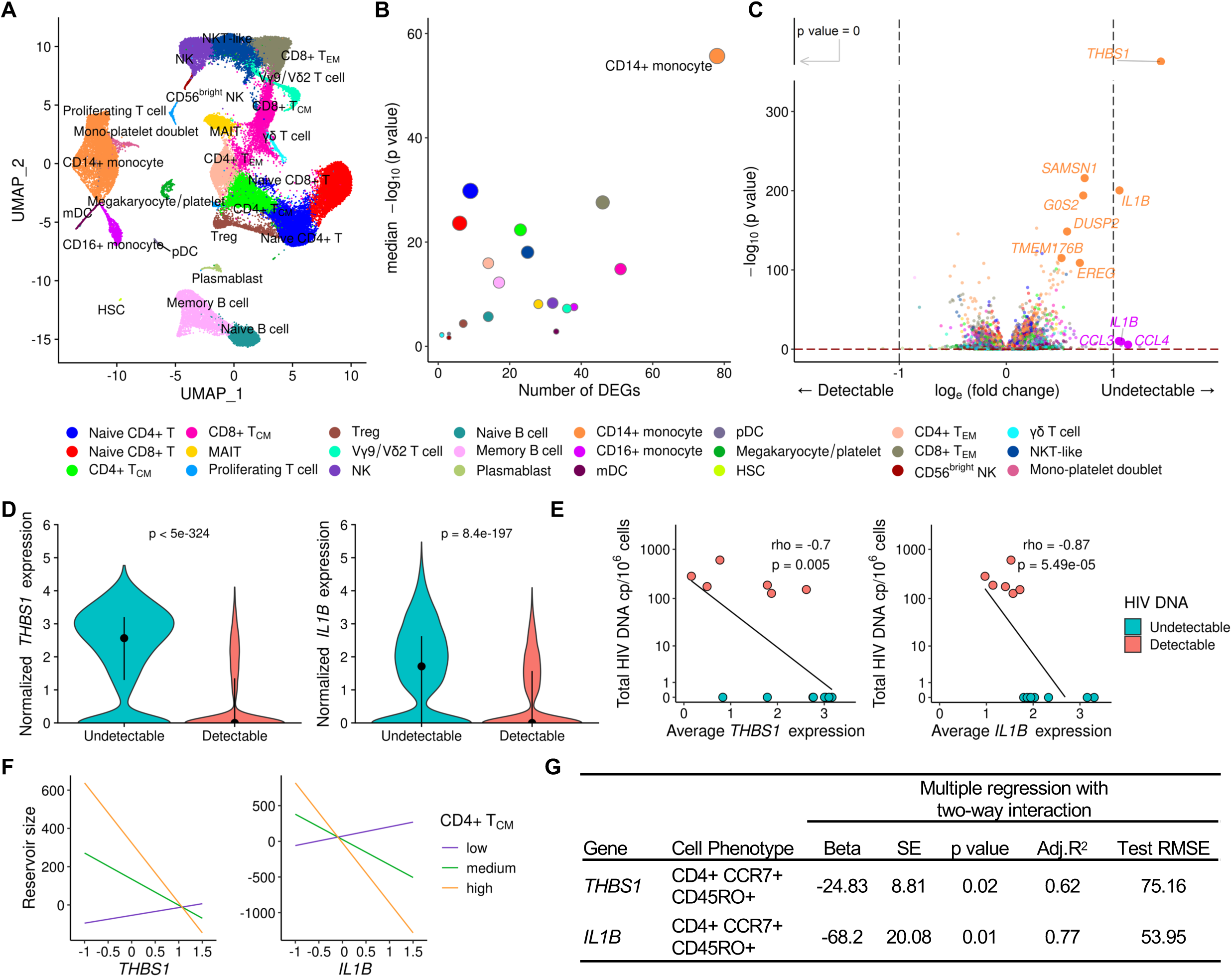
Differentially expressed genes in monocytes associate with HIV reservoir size during ART. A) scRNA-seq identified 24 unique clusters of immune cell subsets. B) CD14+ classical monocytes have the highest number of DEG between the detectable and undetectable reservoir groups. Circle color represents cell subset while circle size indicates corresponding cell number. C) Volcano plot shows DEG in all cell types with p values that are significant after correction, as indicated above the horizontal dotted line. Labeled genes have a p<10e-6 and absolute average log_e_ fold change ≥1 (vertical dotted lines) or p<10e-100 and absolute average log_e_ fold change ≥0.5. D) The most significant DEG in CD14+ monocytes comparing reservoir groups. Black dots represent the median normalized gene expression values (log_e_), and lines represent the interquartile ranges. Teal: undetectable reservoir, red: detectable reservoir. Significance was determined by the Mann-Whitney U test with Bonferroni correction (n=14). E) Participant-specific categorical analyses of the most significant DEGs. Normalized gene expression within CD14+ monocytes was averaged per participant and correlation was determined by the Spearman test (n=14). F-G) Interaction plots of multiple regression between *THBS1 or IL1B* expression in monocytes and reservoir size with varying frequency of the CD4+ T_CM_ population. Nominal p values are indicated for the interaction analyses.

### Monocyte-expressed genes in conjunction with central memory CD4+ T cell frequencies were associated with decreased reservoir size

To understand the association of monocyte gene expression with reservoir size, we used variation in cell frequency data obtained by multi-parameter flow cytometry to determine if specific populations varied between individuals. A total of 117 cell populations were identified and annotated by cell surface marker expression from PBMC isolated at the same time as those used in scRNA-seq analyses. We first used a univariate linear regression analysis of the cell population frequencies and identified CD4+ T cells expressing CD39 on the cell surface as the only marker associated with significantly increased HIV DNA after adjusting for multiple testing (beta=16.9, SE=3.6, P<0.001, q=0.08). In an exploratory analysis, we next evaluated the two-way interaction of each of the 117 phenotypic population frequencies with the top two genes (*THBS1* and *IL1B* in CD14+ monocytes) that associated with decreased HIV persistence. There were several population-specific phenotypic markers whose frequencies increased in the presence of either *THBS1* or *IL1B* and associated with lower reservoir size that were nominally significant (Supplementary table S5). Of the 18 cell populations whose abundances correlated with either the expression of *THBS1* or *IL1B* and associated with decreasing reservoir size, two correlated with both of these genes (Supplementary table S5). The two populations were subsets of central memory CD4+ T cells (CD4+ T_CM_ cells; CD4+CCR7+CD45RO+) that were negative for PD-1 or HLA-DR surface markers (Fig. S5A). Further grouping into other memory CD4+ phenotypes was not possible because of the absence of CD27 surface antibodies in the T cell flow cytometry panel.

However, we observed no differences in frequencies of these CD4+ T_CM_ cell subsets between participants with detectable or undetectable reservoir size (Fig. S5B-D). Given the low frequencies of CD4+ T_CM_ cells that are PD-1+ or HLA-DR+, we combined them with the frequencies of their respective negative populations and obtained their combined parent CD4+ T_CM_ phenotype frequencies. A multiple regression model with two-way interaction also demonstrated a significant inverse relationship of CD4+ T_CM_ frequency and monocyte *THBS1*/*IL1B* expression with reservoir size (P = 0.02 and P = 0.01 for *THBS1* and *IL1B*, respectively, Fig. 2F-G). In this interaction model the significance of *IL1B* over *THBS1* is further strengthened as shown by better accuracy (adjusted coefficient of determination (adj. R^2^)) and deviation (test root mean square error (test RMSE)) metrics (Fig. 2G). Thus, increased expression of *THBS1* or *IL1B* in monocytes in the presence of higher frequencies of CD4+ T_CM_ associated with decreased reservoir size. This suggests that changes in monocyte *THBS1* and *IL1B* expression affect the size of the reservoir via an indirect effect on CD4+ T_CM_, which was the only cell type to show this interaction in PBMC populations measured by flow cytometry.

### Association of *IL1B* expression in monocytes with smaller HIV reservoir size in an independent cohort using different measurements of HIV DNA

To verify the significance of our findings, we used an independent cohort of acutely treated PLWH from the USA (ACTG A5354). This cohort was comprised of 38 male participants of European and African ancestry, with treatment initiated during Fiebig stages III-V. Total HIV DNA reservoir was measured at week 48 after ART initiation. Variation in HIV DNA levels was observed within both the European and African population groups, and scRNA-seq was performed on samples from the week 48 timepoint (Fig. 3A-B). A total of 22 cell subsets were identified (Fig. 3C, Supplementary table S1), the majority of which were consistent with the RV254 cohort from Thailand. In this cohort we expanded scRNA-seq analyses to all available participants with not only detectable or undetectable reservoir, but also the middle group by using HIV DNA measurements as a continuous variable in the MAST statistical framework ^29^. CD14+ monocytes had the greatest number of DEG associating with differences in reservoir size (Fig. 3D). *IL1B* in this single-cell MAST analysis was significant in CD14+ monocytes and validated the directionality seen in the RV254 cohort (coefficient = –0.13, P = 5.1e-34). A categorical analysis of all 38 participants showed an independent significant association of *IL1B* between the extremes as a continuum (p=0.006), but not *THBS1*, with reservoir size (Fig. 3E). Thus, regardless of viral subtype (B or CRF01_AE) or host background (Black, White, Thai), *IL1B* expression in monocytes had a significant inverse association with HIV reservoir size in both the discovery and replication cohorts. Further, IPDA^®^ measurements of the persistent proviruses that comprise the reservoir were available in a subset of the ACTG A5354 study (n=21), where intact or defective proviruses could be analyzed separately. Significantly higher frequencies of persistent intact proviruses compared to defective proviruses after 48 weeks of ART initiation were observed in this cohort of PLWH treated during AHI (Fig. 3F). Participant-specific analyses of *IL1B* expression showed an inverse correlation with different forms of proviruses (Fig. 3G). Notably, when we harnessed the power of single-cells using the MAST framework for continuous analyses, the inverse *IL1B* association was significant across most forms of the persistent provirus as measured by IPDA^®^, showing that findings were valid across total, intact and defective proviruses (Fig. 3G).

**Figure 3.**
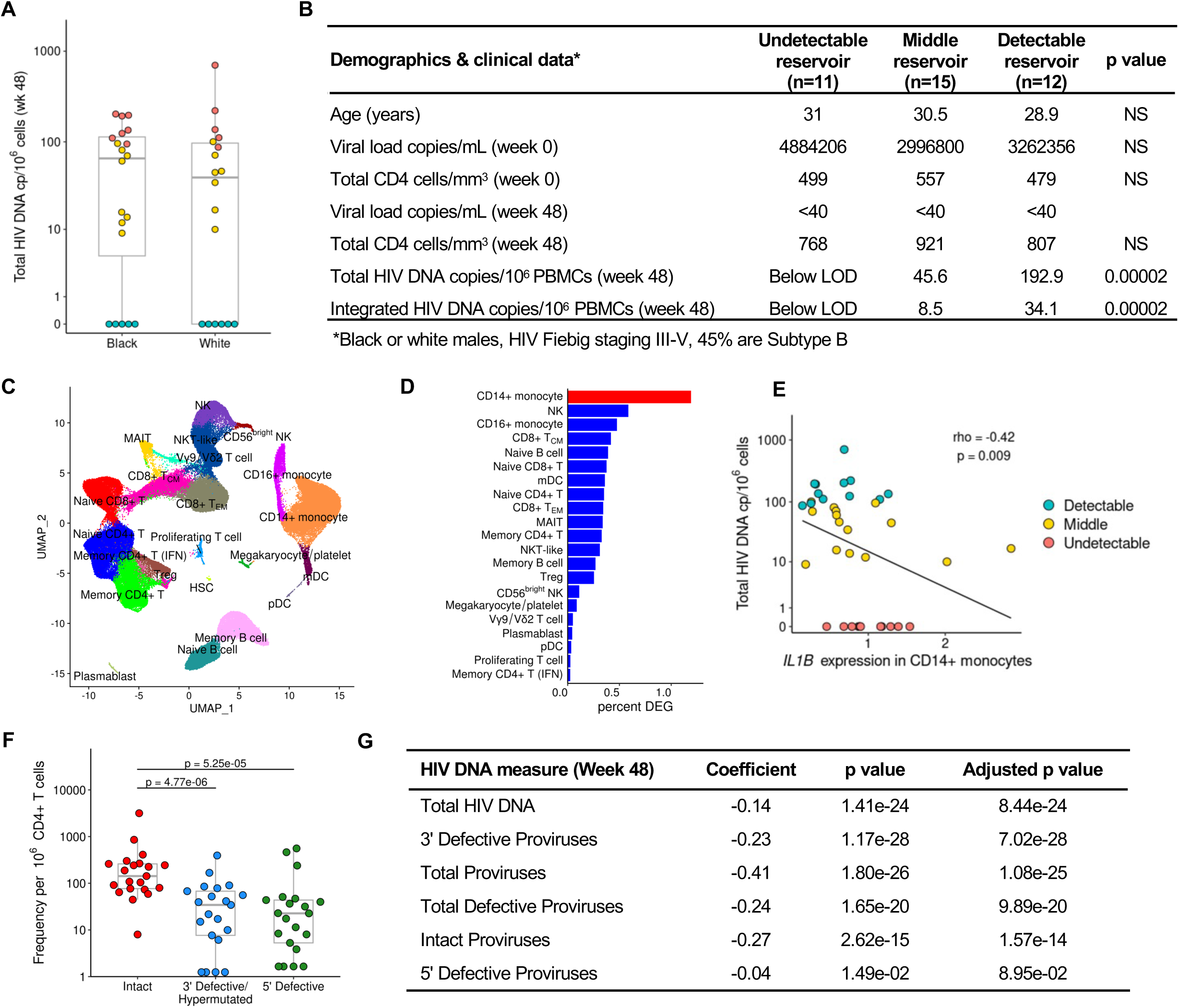
Validation of *IL1B* association with smaller reservoir size from an independent cohort with a different infecting viral subtype across various Fiebig stages. A) HIV DNA levels vary within the A5354 subtype B cohort from the USA (n=38). The participant samples used in this study are highlighted based on reservoir size: red=detectable; teal=undetectable and yellow=middle. Black and White indicate differences in ancestry of the participants. B) Characteristics of participants comprising the detectable, middle, and undetectable reservoir size categories (mean values are shown) and p values comparing the extreme phenotype groups. HIV-1 subtype information was only available for a subset of the participants. NS: not significant C) Dimensionality reduction plot of the different immune clusters in this cohort. D) CD14+ monocytes have the highest number of normalized DEG associated with reservoir size using a continuous analysis including all 38 participants. E) *IL1B* participant-specific average gene expression in CD14+ monocytes categorized by total HIV DNA (n=38). Spearman correlation p value and rho are shown. F) IPDA® measurements from a subset of the participants in this cohort (n=21). G) *IL1B* association with different reservoir type measurements (rows) from the participants with IPDA measurements (n=21).

### Transcriptional programs implicated NF-κB with the differences in HIV-1 reservoir size

Given these significant effects of individual monocyte genes on reservoir size, we explored the broader consequences of transcriptional changes in monocytes using unbiased weighted gene co-expression network analyses (WGCNA) and identified nine modules of co-expressed genes within CD14+ monocytes from RV254 (Fig. 4A). The second largest module, M3, was significantly more highly expressed in the original Thai cohort from participants with an undetectable reservoir and contained 452 genes, including *IL1B* (P_adjusted_<5e-324, Fig. 4B). Comparing expression between the detectable and undetectable groups in the independent A5354 cohort using the M3 module genes identified in the Thai cohort, we confirmed that this module was similarly enriched in the cells from the undetectable group (P_adjusted_= 1.3e-55, Fig. 4C). There were no other modules that were significantly associated with the reservoir size in both studies. Expression of the top 25 hub genes in this M3 module was generally higher in the undetectable than in the detectable reservoir group in both cohorts (Fig. 4D). The strength of this signature was further reinforced by the predicted interaction of the genes at the protein level (Fig. 4E). Gene ontology analyses showed that genes in the M3 module were enriched in several pathways, including regulation of apoptosis and NF-kappa B (NF-κB)(Fig. 4F). TNFα signaling via NF-κB had the largest membership of genes from the M3 module. Complementing the findings in CD14+ monocytes, once again the same signaling pathway was also enriched in a module that was highly expressed in the undetectable reservoir group in the memory CD4+ T cell subset, suggesting an effect on NF-κB signaling in the cell population which harbors the latent reservoir (Fig. S6). These pathways are consistent with the *IL1B* findings, suggest a broader change in the inflammatory homeostatic state, and may define a coordinated transcriptional change that accompanies *IL1B* expression differences which associate with reservoir size.

**Figure 4.**
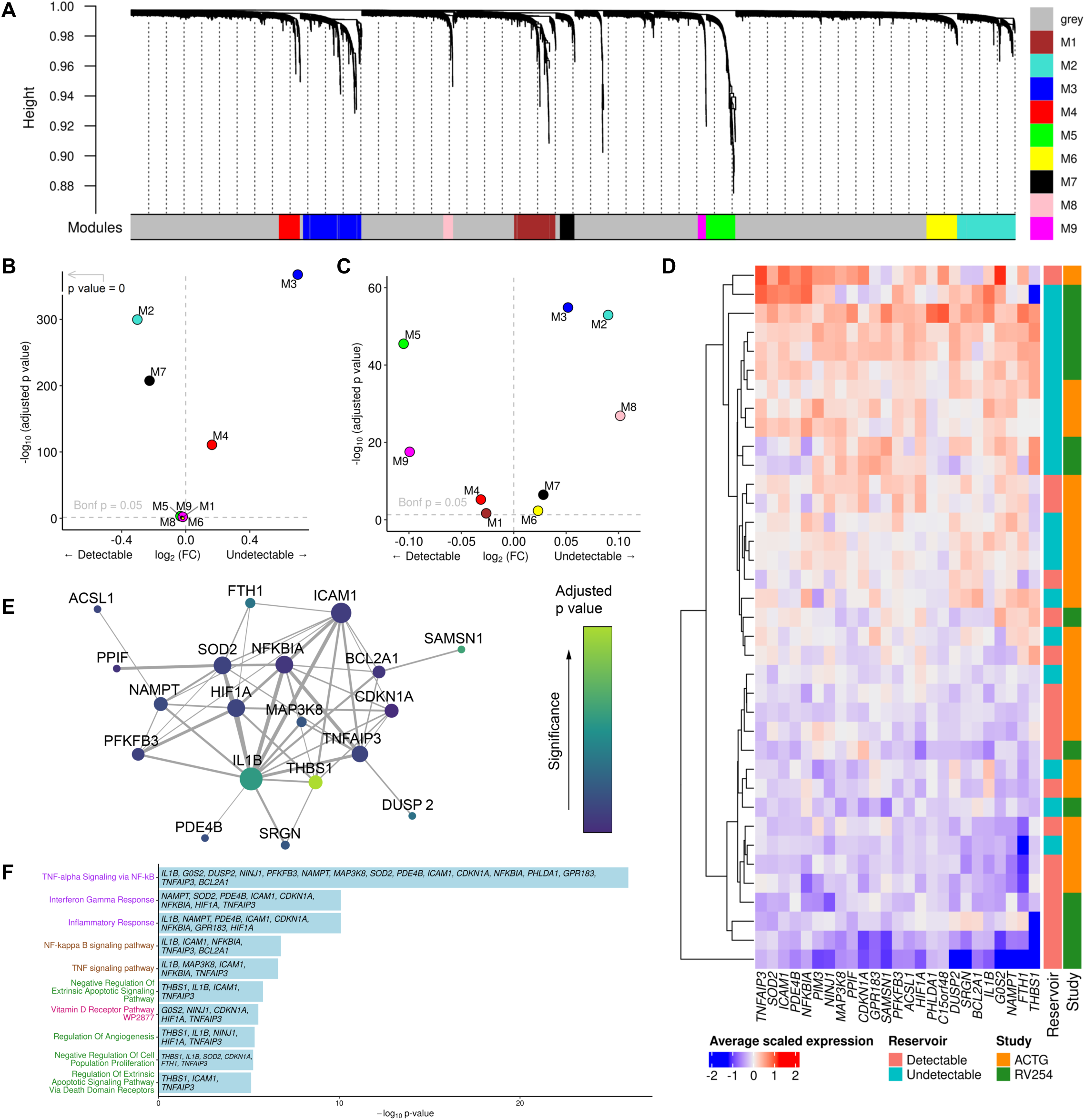
Pathway analyses identifies a distinct signature associating with reservoir size. A) Gene co-expression modules in CD14+ monocytes from the RV254 Thai study. B) *IL1B* is in the M3 WGCNA module which was enriched in cells from RV254 participants with undetectable reservoir based on the top 25 hub genes in the module (detectable=6, undetectable=8). C) Using the same module hub genes found in RV254, the M3 module was also enriched in cells from the undetectable reservoir participants in the A5354 cohort when HIV DNA levels were grouped categorically (detectable=12, undetectable=11). D) Average expression of the 25 top hub genes from the M3 module had generally higher expression in participants with undetectable reservoir in both cohorts. E) Predicted protein interaction network of top 25 hub genes using the STRING protein database. Larger nodes have higher degree of connectivity; node color indicates significance in the categorical DEG comparison between the detectable and undetectable groups in RV254 CD14+ monocytes. F) Gene ontology analyses of genes enriched in module M3 in CD14+ monocytes.

### IL1B activates NF-κB, enhancing productive HIV infection while inhibiting viral spread in vitro

Binding of IL1B to its IL1 receptor induces a signaling cascade ultimately leading to the activation of NF-κB ^30^. This transcription factor plays a key role in LTR-mediated transcription of proviral DNA, and its stimulation is well known to reactivate latent HIV-1 ^31,32^. Thus, we explored whether activation of NF-κB in CD4+ T cells could explain why increased monocyte *IL1B* expression could reduce the size of the latent HIV reservoir. To assess whether IL1B activates NF-κB, we treated A549 NF-κB reporter cells, which express the secreted embryonic alkaline phosphatase (SEAP) reporter gene under the control of the IFN-β minimal promoter fused to five NF-κB binding sites, with IL1B, TNFα, LPS or medium only. We observed that IL1B increased NF-κB activity ∼4-fold regardless of HIV-1 infection status (Fig. 5A). Next, we examined the ability of IL1B to activate NF-κB in primary human T cells. Degradation of inhibitory IκB proteins is a critical step in the activation of NF-κB, and their phosphorylation one of the earliest and most specific events in this process. Treatment with IL1B increased IκB phosphorylation and strongly reduced the overall levels of inhibitory IκB indicating activation of NF-κB (Fig. 5B).

**Figure 5.**
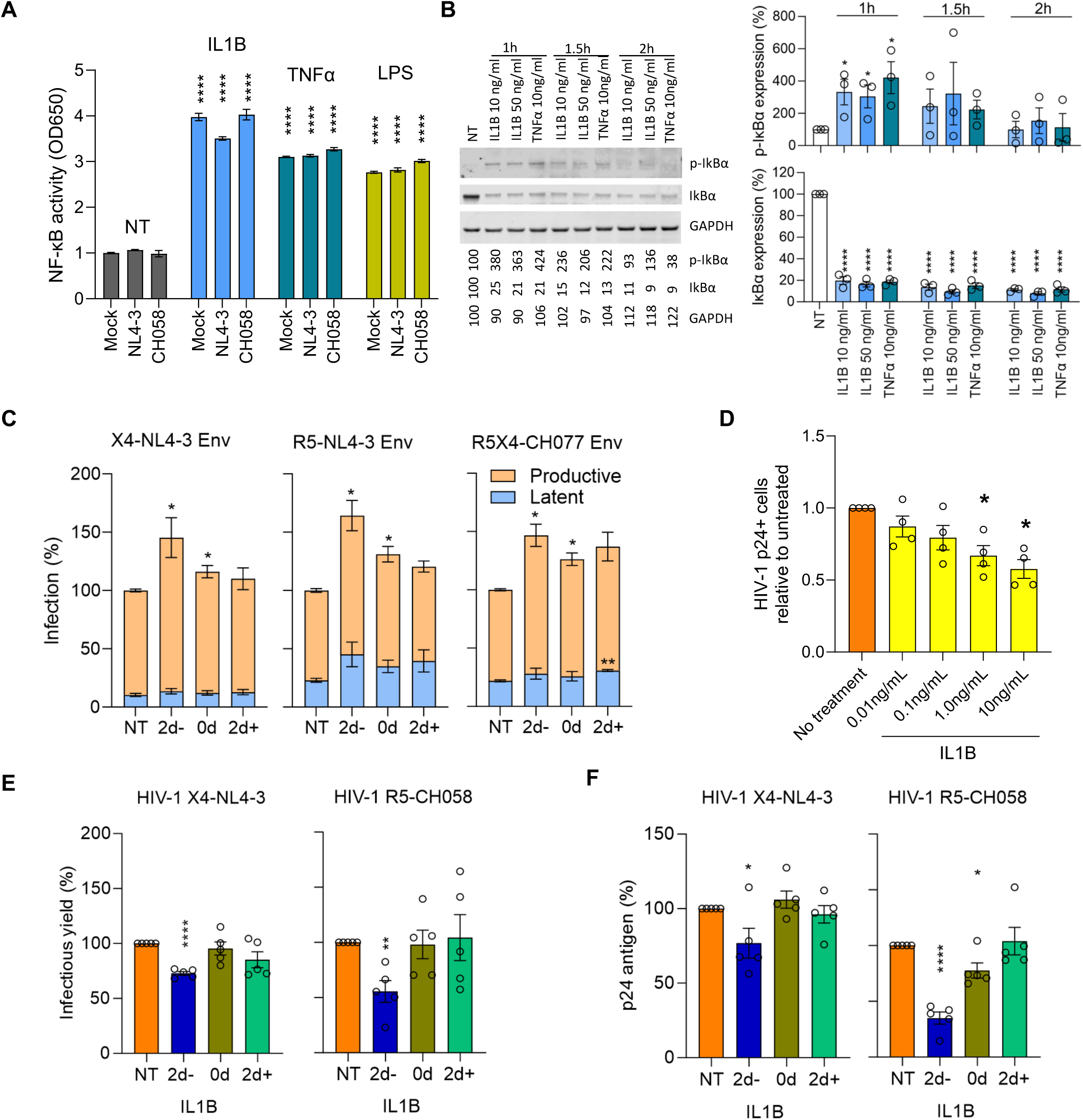
*In vitro* IL1B activates NF-κB, increases HIV proviral transcription, and inhibits spreading infection. A) Effects of IL1B on NF-κB activity were assessed using A549 NF-κB reporter cells. Cultures were treated with IL1B, LPS, and TNFα and infected with VSV-pseudotyped NL4-3, CH058, or Mock control. After 24h the Alkaline Phosphatase Blue Microwell assay was performed with OD650 values relative to no treatment control (NT) reflecting NF-κB expression which is shown on the Y-axis. B) PBMC from 3 donors were treated with IL1B or TNFα and examined for IκBα phosphorylation as described in the methods section. Graphs present the protein expression from these donors; unpaired t test, *p<0.05, ****p<0.0001. C) Effects of IL1B in vitro when HIV was quantified after a single round of infection. Plots show the relative proportions of pMorpheus-V5 latently (blue) or productively (orange) infected PBMC in cultures treated with IL1B prior to, simultaneously, or after transduction with Env viral particles carrying the indicated Env protein. The data represent the average of 3 individual healthy donors, with error bars representing the average ±SEM, and statistical significance was established using unpaired t tests; *p<0.05, **p<0.002. D) Effects of IL1B on spreading HIV-1 infection in cell culture. Using HIV-1 YU-2, bar plots display the relative p24-positive cell fractions after pre-treatment with increasing concentrations of IL1B (from 0.01-10.0 ng/mL, 10-fold increments) across four different donors. E, F) Bar plots display the average infectious virus yields (E) and p24 antigen levels (F) at 4 days post-infection relative to the no IL1B treatment controls normalized to 100%; unpaired t test, *p<0.05, **p<0.002, ***p<0.0002. Corresponding replication curves are shown in Fig. S8.

To more directly determine the impact of IL1B on latent and productive HIV-1 infection, we used the HIV molecular clone pMorpheus-V5, which lacks a functional *env* gene but encodes all accessory proteins ^33^. Cells productively infected with pMorpheus-V5 express V5-NGFR (Nerve Growth Factor Receptor) driven by the PGK (phosphoglycerate kinase) promoter, as well as HSA and mCherry driven by HIV-LTR, while latently infected cells express only V5-NGFR. Activated PBMC from three healthy donors were treated with IL1B prior to, simultaneously, or after transduction with pMorpheus-V5 pseudo-typed with the Env proteins of WT X4-tropic NL4-3, an R5-tropic derivative thereof ^34^, and the dual R5X4-tropic CH078 transmitter-founder strain ^35^. The frequencies of latently infected cells were generally lower compared to productive infection and in most cases not significantly altered by IL1B treatment. In contrast, the number of productively infected cells increased significantly (P < 0.05) for all 3 viruses when exposed to IL1B treatment during or 2 days prior to infection (Fig. 5C).

NF-κB plays a complex role in HIV infection because it not only activates LTR transcription but also plays a key role in innate immunity and induces expression of numerous antiviral factors ^36,37^. Indeed, pretreatment of stimulated PBMC with IL1B for two days resulted in a dose-dependent decrease in infection with the R5-tropic YU-2 virus as determined by the frequencies of sorted p24+ CD4+ T cells (Fig. 5D, S7A). To further explore the effect of IL1B on spreading HIV-1 infection, stimulated PBMC from five donors were treated with IL1B prior to, at the same time, or after infection with HIV-1 NL4-3 or the transmitted founder CH058 molecular clone. Infectious virus production at 2, 4, 6 and 9 days post-infection was determined by p24 ELISA and infection of TZM-bl indicator cells. Infectious virus yields peaked at day 4 in most of the infected PBMC cultures (Fig. S8). Two-day pretreatment with IL1B generally reduced viral replication compared to the untreated controls. Both infectious virus and p24 antigen production by NL4-3 and (more strongly) the primary CH058 strain were significantly (P < 0.05-0.001) reduced at 4 days post-infection (Figs. 5E, 5F, S8). In comparison, only modest effects were observed when IL1B was added during or after infection, presumably because the induction of an antiviral state requires de novo synthesis of antiviral factors. Notably, the levels of cell death were low and did not differ significantly from the uninfected control (Fig. S9).

IL1B is also known to affect the differentiation of CD4+ T cells into various subsets ^38^ that may differ in their susceptibility to HIV-1 infection. Notably, in the YU-2 infected cultures over 95% of p24+ cells expressed the CD45RO memory T cell marker, consistent with previous reports ^39,40^, and p24+ populations exhibited higher frequencies of both CD4+ T effector memory and transitional memory subsets (CD4+ T_EM_ and CD4+ T_TM_, respectively) compared to the p24-populations (Fig. S7B-C). These findings underscore the importance of subset phenotypes for HIV infection and suggest that IL1B could alter the frequencies or phenotypes of HIV-susceptible CD4+ T cell subsets to modulate HIV reservoir size. We observed an IL1B dose-dependent increase in the frequency of CD4+ T_CM_, accompanied by decreases in the frequencies of both CD4+ T_TM_ and CD4+ T_EM_, when PBMC from healthy participants were cultured in vitro (Figs. S10A-B). Altogether, our in vitro data suggest that IL1B could decrease the HIV reservoir size in vivo through multiple mechanisms, including by promoting NF-κB mediated activation of latent HIV, inducing innate antiviral factors and changing the composition of T cell populations (Fig S11).

## DISCUSSION

In this study, we used an unbiased high throughput single-cell approach to identify differences in host transcriptional profiles that associate with the size of the viral reservoir in acutely treated PLWH by screening extreme phenotypes of reservoir size. Recent single-cell transcriptomic studies have focused on the effects of differentially expressed host genes specifically in CD4+ T cells from PLWH on treatment ^20,41^, but it is important to also examine other cell populations that might influence the viral reservoir. We observed significant differences in the gene expression profiles of multiple immune cell subsets even after almost one year of complete viral suppression on ART, distinguishing participants with variably-sized viral reservoirs, which were conserved across two cohorts comprising a total of 52 individuals and encompassing multiple host and viral genotypes. Significant differences were discovered in part due to accounting for potential confounders by selecting participants matched for Fiebig stage at the time of ART initiation, viral subtype, and sex prior to examining gene expression in single cells from participants in the Thai discovery cohort. These differences were generalizable to a subtype B cohort comprised of participants with greater variability and having IPDA® reservoir measurements. Frequencies of defective viruses in the ACTG study were lower than intact proviruses when measured by IPDA® which is not surprising considering the timing of sampling after ART initiation ^42–46^. Regardless, the single-cell association of *IL1B* with reservoir size remained significant with different reservoir measures. These findings were also enabled by the use of scRNA-seq with its advantage compared to bulk transcriptomics that gene expression differences can be traced to specific cells rather than to “averaged” signals from heterogeneous populations.

We found that monocytes, specifically the CD14+ subset, showed the highest number of enriched pathways and DEGs, with *IL1B* being associated with differing reservoir sizes in two independent cohorts using various measurements of total, defective, and intact proviruses. IL1B is a potent proinflammatory cytokine ^47^, expressed in cells such as monocytes, neutrophils, B cells, and DCs, that is involved in a variety of cellular activities. Though *IL1B* was the most significant DEG, we also detected a network of coexpressed genes that support a coordinated change in the monocyte transcriptional profile. Together, these findings are consistent with our previous observations that monocytes can play an important role both after vaccination and after treatment initiation when virus is suppressed ^48^. In addition to assessing differences in host gene expression, frequencies of all major cell populations comprising PBMC were assessed by surface protein-based flow cytometry. These frequencies were evaluated in the context of scRNA-seq gene expression for potential combined effects on peripheral blood reservoir size. Of the 117 immune population frequencies assessed by flow cytometry, only CD4+ T_CM_ and its subsets were associated with a smaller reservoir size when *IL1B* expression levels in CD14+ monocytes were high. Interestingly, this is supported orthogonally by a recent report that CD4+ T_CM_ host the smallest quantity of intact proviruses compared to naïve and other memory subsets ^7,49^. Our findings suggest a link between DEG in monocytes from different extreme reservoir size phenotypes and a specific CD4+ T cell subset previously implicated as harboring the latent HIV reservoir.

How exactly IL1B expression levels in monocytes influence the reservoir in vivo remains to be determined. However, IL1B-mediated activation of NF-κB is well established ^30,50^ and may explain the link between increased *IL1B* expression in CD14+ monocytes and a reduced latent HIV reservoir size (Fig. S11). Besides IL1B, TNFα is one of the strongest endogenous inducers of NF-κB and the pathway “TNFα signaling via NF-κB” had the largest number of enriched genes in the M3 and M4 modules in CD14+ monocytes and memory CD4+ T cells, respectively. The key role of NF-κB in proviral HIV-1 gene expression has been known for decades. However, NF-κB also plays key roles in immunity and inflammation, inducing numerous antiviral factors ^36,37^. Notably, NF-κB activates LTR transcription directly, while inhibitory effects require de novo synthesis of antiviral factors. Thus, HIV-1 and lentiviruses tightly regulate NF-κB activity to enable viral transcription while minimizing antiviral gene expression ^51–53^. The induction of innate antiviral immunity by NF-κB may reduce viral reservoir seeding during acute infection. However, induction of proviral transcription by NF-κB is likely the more important mechanism in ART treated individuals, where viral replication is effectively suppressed and induction of productive infection renders the latent reservoir susceptible to elimination. IL1B, TNFα and NF-κB all play complex roles in the survival, activation, and differentiation of T cells and other immune cells ^38^. Thus, they may also impact the frequency of reservoirs harboring cells by more indirect mechanisms, such as shifts in the T cell subtype composition or cell survival. IL1B is best known for its role as a secreted cytokine. In some cases, however, it may also act in a cell-associated manner and the potential of IL1B-expressing CD14+ monocytes warrants further investigation. Notably, latency reversing agents that stimulate NF-κB have been extensively studied in shock-and-kill approaches and shown to reactivate HIV-1 from latency in both CD4+ T cell latency models and HIV-1-infected patient-derived cells ^54–56^. Thus, it is tempting to speculate that, similar to TNFα, IL1B acts as a natural NF-κB inducing latency-reversing agent.

In addition to the strongest effect observed of higher *IL1B* levels, we also detected that *THBS1* in CD14+ monocytes associated with a smaller reservoir of infected cells. The association of *THBS1*, encoding for thrombospondin, with reservoir size was only observed in the RV254 Thai cohort, and not validated in A5354 participants from the USA, most likely attributable to differences in genetic ancestry and so our mechanistic analyses focused on *IL1B*. However, population specific associations are important and further studies are warranted to understand the effect of *THBS1* on reservoir size given that anti-HIV properties have been attributed previously ^57^. Similarly these studies were limited to AHI cohorts and exploration in the context of untreated chronic infection is also necessary. Another limitation to this unbiased single-cell approach is that it lacks equal power to detect effects across all cellular subsets and could miss effects in populations less frequent than monocytes. Additionally, how other monocyte genes or additional factors interact in vivo to influence the decreased reservoir size observed in participants with increased monocyte *IL1B* remains to be investigated further. Though intact proviral DNA measurements are still being adapted for use beyond subtype B ^42^ and therefore were not available for participants in the RV254 cohort, we were able to bridge our findings by leveraging IPDA® measurements of persistent intact and defective proviruses in the ACTG A5354 cohort. Although the strength of the associations differed, *IL1B* remained inversely associated with smaller reservoir size regardless of proviral intactness. It is plausible that different host factors exert effects depending on the form of the provirus and raises the need for additional in-depth investigations.

Overall, our findings support that immune cells other than T cells can modulate the HIV reservoir in a clinical cohort, and that this effect may be influenced by specific genes and pathways. These findings were based on unbiased single-cell approaches and give rise to new hypothesis-driven questions that should be tested in other cohorts, where further confirmation of host cellular gene products and pathways involved in reservoir formation or maintenance may provide targets for therapeutic intervention and remission strategies. In particular, our implementation of single-cell approaches that are unbiased and allow broad screening for effects that impact reservoir size in ART treated individuals in vivo suggests that IL1B-induced NF-κB dependent mechanisms for induction of proviral transcription may represent latency reversing strategies which could be effective in vivo.

## MATERIALS AND METHODS

### Study design

Demographic and clinical data (viral load, CD4+ T cell counts, HIV DNA, HIV subtype, Fiebig stage) were available from 163 acutely-treated PLWH from the men who have sex with men (MSM) RV254/SEARCH010 cohort (NCT00796146) in Thailand ^13,24,58^. For discovery analyses, scRNA-seq, immune receptor sequencing (TCR and BCR), and flow cytometry were performed on initially cryopreserved PBMC from 14 selected participants in this cohort and validated in an additional 38 male participants from the ACTG A5354 study (NCT02859558), a single-arm, open-label study to evaluate the impact of ART initiation during AHI conducted at 30 sites in the Americas, Africa, and Asia. Samples from all participants were collected at 48 weeks post-ART initiation. In a subset of participants samples were also available from week 0 at AHI. Blood from healthy participants without HIV for in vitro experiments was obtained from the WRAIR 2567.05 protocol. All participants provided informed consent, and use of samples for research was approved by institutional review boards in Thailand and the USA.

### HIV Reservoir measurements

Total HIV DNA and integrated DNA were measured by quantitative PCR (qPCR) as described previously ^14,28^. Briefly, pellets of PBMC or CD4+ T populations were suspended in 15ul of Proteinase K lysis buffer per approximately 100,000 cells, and digested for 18h at 55°C. Total HIV DNA was quantified using primers and a probe situated in the 5’-LTR, while primers and probe used for integrated DNA were situated in Alu and the 5’-LTR. ACH-2 cells, which carry a single copy of the integrated HIV genome, were used to generate a standard curve for both assays. The cell input for each of the three replicates was approximately 100,000 per replicate (∼300,000 total) and the lower limit of detection of this assay was 3.3 copies/10^6^ cells. Participants were grouped into detectable or undetectable reservoir based on the presence of total HIV and integrated viral DNA measured independently in both PBMC and CD4+ T cell populations, depending on sample availability for the latter. Presence of integrated HIV DNA was used as a criterion for defining the categorical groupings. The reservoir phenotype was defined as undetectable when both total and integrated HIV DNA were below the LOD. In contrast, the reservoir was defined as detectable when total HIV DNA > LOD.

### IPDA

Accelevir Diagnostics performed HIV-1 intact proviral DNA assays (IPDA®) to discriminate between, and separately quantify, the frequencies of intact and defective persistent proviruses. The design and performance of this assay have been described previously ^42,59^. Briefly, cryopreserved PBMC were thawed and CD4+ T cells were isolated and assessed for cell count, viability, and purity by flow cytometry. RNA-free genomic DNA was then isolated from the recovered CD4+ T cells, with concentration and quality determined by fluorometry and ultraviolet-visible (UV/VIS) spectrophotometry, respectively. The IPDA® was performed, and data reported as proviral frequencies per million input CD4+ T cells. These procedures were performed by blinded operators using standard operating procedures.

### Single-cell RNA library generation and sequencing

PBMC from the 14 RV254 participants on ART for 48 weeks were washed, resuspended in PBS plus 0.5% FBS, and simultaneously processed for scRNA-seq and flow cytometry. A total of 50,000 cells (at 1,000 cells/ul) from each donor were set aside for scRNA-seq library construction and the remaining cells were used for flow staining as described later. The diluted PBMC suspensions were prepared for scRNA-seq using the Chromium Next GEM 5’ Single Cell V(D)J Reagent Kit and the Chromium Controller (both 10x Genomics) per manufacturer’s instructions. Briefly, targeting a recovery of 8,000 cells/donor, samples were loaded into separate wells of Chromium chips. Amplified cDNA was used to make gene expression (GEX), TCR, and BCR libraries. The GEX library construction used a 14 or 16 cycle Sample Index PCR program, based on amplified cDNA concentrations. PBMC from the 38 A5354 participants were individually stained with TotalSeq-C anti-human hashtag antibodies (BioLegend), batched, and processed for gene expression (GEX) and hashtag oligo (HTO) libraries as previously described to improve cost-effectiveness ^26^. Cells from each batch were loaded into 4 different wells of Chromium chips for targeted recoveries of 16,000 cells/well.

Libraries from both studies were then assessed for quality and concentrations using the DNA High Sensitivity D5000 ScreenTape Assay with the TapeStation (both Agilent, CA), and subsequently pooled and quantitated with a MiSeq Nano Reagent Kit v2 (300 cycles) (Illumina, CA) sequencing run. Final libraries were sequenced using the NovaSeq 6000 S4 Reagent Kit (300 cycles) on a NovaSeq 6000 instrument (both Illumina).

### Multiparameter flow cytometry

PBMC from 14 participants were stained with Aqua Live/Dead stain (Molecular Probes), washed, and blocked using normal mouse IgG (Caltag). The cells from each donor were then split into four to run four different polychromatic flow panels using conjugated fluorescently labeled monoclonal antibodies against several surface markers to define B, T, Myeloid, and NK cell subsets (Extended table 2). For the T cells panel, cells were pre-stained with an MR-1 tetramer ^60^ prior to staining for additional surface markers. Following surface marker staining, cells were washed, permeabilized and fixed with FoxP3 Fixation/Permeabilization Set (eBiosciences). Cells were then washed, stained intracellularly, washed again, and analyzed using a BD FACS Symphony A5. Data were analyzed with FlowJo v.9.9.6 or higher (Becton Dickinson).

### Virus production

HEK293T cells were transfected with NL4-3, CH058 and pMorpheus proviral constructs using the TransIT-LT1 (Mirus) transfection reagent per manufacturer’s protocol. Infectious molecular clones of HIV CH058 and CH077 were kindly provided by Beatrice H. Hahn ^52,61^. Media were changed 24h post transfection and virus stocks were collected 24h later. PBMC were infected with freshly produced virus. The HIV YU-2 infectious molecular clone stock was obtained from the HIV Reagent Program.

### In vitro functional characterization

#### Effects of IL1B on cell population frequencies and HIV infection

PBMC were isolated from the blood of healthy donors by density centrifugation on a Ficoll-Paque gradient (GE Healthcare) and stimulated by anti-CD3/CD28 Dynabeads at a 1:1 ratio with the estimated CD4+ T cell population in PBMC (25% in total PBMC) in Complete Cell Culture Medium (RPMI Medium, GlutaMAX supplemented HEPES with 10% fetal bovine serum, penicillin/streptomycin) (all Gibco) supplemented with 40 U/ml IL2 and with or without recombinant IL1B at four different concentrations (0.01-10 ng/ml, at 10-fold intervals) (both R&D Systems) for 4 days. Treated PBMC were either immediately analyzed by flow cytometry to assess frequencies of T cell subpopulations, or infected with an R5 tropic molecular clone, YU-2, at a concentration of 1 µg of p24 per million cells and cultured for a further two days before assessing the relative frequencies of infected cells by flow cytometry.

#### Effects of IL1B on HIV infectivity

PBMC isolated from healthy donors were treated with 10 ng/ml IL1B concentrations at different times relative to HIV infection initiation: pre-treated 2 days prior to infection, added simultaneously, or added 2 days post-infection. Briefly, PBMC were isolated by Ficoll gradient centrifugation and cells were stimulated by PHA in Complete Cell Culture Medium with 100 U/ml IL2 for 3 days. On day 1 post-isolation the required cells were set aside for IL1B treatment for 2 days prior to infections. On day 3 post-isolation, PBMC were infected by spinoculation (1200 xg, 2 h, 26°C) with 150 ng of freshly produced NL4-3, or 500 ng of CH058 virus strains, per million cells. After spinoculation cells were washed 5 times with 1x PBS and resuspended in fresh medium containing IL2 and IL1B per the schedule. Cells were cultured for an additional 9 days during which 400 µl of supernatant was removed every second day for determination of infectious virus yields. To determine infectious virus yield, 10,000 TZM-bl reporter cells per well were seeded in 96-well plate. The next day cells were infected in triplicate for 9 days with the collected supernatants. Three days post infection the TZM-bl cells were lysed and *b-galactosidase* reporter gene expression was assessed with the GalScreen Kit (Applied Bioscience) per manufacturer’s protocol using an Orion microplate luminometer (Berthold).

### Flow cytometry staining of pMorpheus infection

PBMC infected with pMorpheus were collected on day 5 post-infection and stained for membrane marker V5 and viability. Cells were harvested, washed 3x with PBS and resuspended in surface staining of V5 and viability dyes (V5 Alexa Fluor 647; eBioscience fixable viability dye efluor 780; both Thermo Fisher Scientific). After 30 min incubation, cells were washed 3x with PBS, fixed with 4% PFA for 30 min and analyzed by flow cytometry.

### Western blotting

PBMC were isolated as described and cultured for 3 days with IL2 and PHA. On day 3 cells were treated with IL1B (10 and 50 ng/ml) and TNFα (10 ng/ml) for 1, 1.5, and 2 hr. Cell lysates were prepared and western blotting was performed as described previously ^62^. Proteins were stained with the following primary antibodies: phospho-IκBα (Cell Signaling), IκBα (Santa Cruz), GAPDH (BioLegend).

### NF-κB reporter assay

A549-Dual^TM^ Cells (InvivoGen) were seeded at a density of 20,000 cells per well on 96-well plates. Cells were treated on the following day with IL1B (10 ng/ml), TNFα (10 ng/ml), LPS (1000 U/ml) and infected or not with VSV-G pseudotyped HIV-1 NL4-3 or CH058 for 24h when the Quanti Blue assay was performed as described by the manufacturer (Invitrogen).

### Viability assay

Cells were harvested, washed once with PBS and stained for 15 min at RT in the dark with eBioscience fixable viability dye 780 (Thermo Fisher Scientific). Cells were then washed twice with PBS, fixed in 2% PFA for 30 min at 4 °C and analyzed by FACs.

### Bioinformatics analyses

#### Sequence data processing

Single-cell gene expression data from PBMC were generated using the 10x Genomics Cell Ranger pipeline (v3.0.0 – 3.1.0) (10x Genomics) per manufacturer’s recommendations and the 10x Genomics human reference library (GRCh38 and Ensembl GTF v93). For the RV254 sequencing run without hashing, the average number of genes per cell was 1,236 and the average number of unique molecular identifiers (UMI) was 3,288. The mean read depth per cell was approximately 103,000-236,000 reads. The minimum fraction of reads mapped to the genome was 92.95% and sequencing saturation was on average greater than 94%. For the hashed A5354 sequencing runs, the average number of genes per cell was 1,432 and the average number of UMIs for RNA transcripts was 4,192. The mean read depth per cell was approximately 69,000-88,000 reads for the gene expression library and 9,000-14,000 reads for the antibody library. The minimum fraction of gene expression reads mapped to the genome was 88.5% and RNA sequencing saturation was on average greater than 89%. Downstream analysis of Cell Ranger outputs including quality filtering, normalization, multi-sample integration, visualization, and DEG were performed using the R package Seurat (v3.1.1 – 4.3.0).

#### RV254 gene expression processing

Cells with mitochondrial percentages greater than 10% and cells that had <200 or >6,000 expressed genes were removed from analyses. 62,925 cells remained after the quality control (QC) process. After log-normalization with a scale factor of 10,000, the top 2000 variable features within each sample were selected. We found integration anchors using dimensions 1:30 and integrated cells from all 14 participants. Shared Nearest Neighbor-based (SNN) clustering was performed using the top 30 principal components (PC) with a resolution of 0.5, and cells in the clusters were visualized by UMAP projection. Cluster marker genes were determined using Seurat FindAllMarkers and cluster identities were manually annotated using differentially expressed genes between the clusters and known lineage cell markers.

#### A5354 demultiplexing and gene expression processing

HTO expression matrices were normalized, demultiplexed, and assigned to specific participants using the methods described ^26^. Negative cells and cells with greater than 10% mitochondrial gene expression were removed. Gene expression matrices (containing a total of 21,870 genes) for all 38 participants and for doublet cells were normalized. We performed reference-based integration using two participants from each ADT batch as the references. Cells that were identified as doublets via hash demultiplexing and cells in clusters from an initial round of QC that were enriched for doublets or had high expression of *HBB* were removed and SNN clustering at resolution 0.3 was performed on the remaining 140,172 cells. Clusters were visualized and annotated using lineage markers and differentially expressed genes similar to the process for RV254. No γδ T cell or monocyte-platelet aggregation clusters were identified, and CD4+ memory T cells were comprised of one large cluster and one smaller cluster with upregulation of interferon-induced genes, instead of subsets of CD4+ T cells as observed in RV254.

#### Differential gene expression

Categorical differential gene expression analyses within each cell type subset between the two reservoir groups was performed within Seurat using a Mann-Whitney U test with Bonferroni correction (n=19,581). Genes that were not expressed in at least 10% of cells in either group or that did not have a log fold change of >|0.25| were excluded from consideration, as were mitochondrial and ribosomal protein genes. The MAST framework was implemented to examine correlation of gene expression of different cell subsets with the continuous total HIV DNA measurements as the outcome ^29^. Genes with expression frequencies <10% were removed before analyses. Results from each cell subset were corrected for multiple testing using the Bonferroni correction. Genes without a beta coefficient >|0.1| and additional manually curated genes were excluded from consideration. Continuous MAST analyses for a subset of 21 participants with IPDA® data was performed to see if *IL1B* remained significant using different reservoir measurement parameters.

Participant-specific expression values were generated for certain genes using Seurat’s AverageExpression in CD14+ monocytes within participants on the log-normalized expression data. <u>TCR/BCR sequence analyses:</u> TCR/BCR clonotype identification, alignment, and annotation were performed using the 10x Genomics Cell Ranger pipeline (v6.1.2; 10x Genomics) per manufacturer’s recommendations. Clonotype alignment was performed against the Cell Ranger human V(D)J reference library 7.1.0 (GRCh38 and Ensembl GTF v94). The Cell Ranger clonotype assignments were used for both BCR and TCR Clonotype visualization and diversity assessments, and analyses were performed using R for IG chains within annotated B cell types (memory B cells, naïve B cells) or TRA/TRB chains within annotated T cell types (CD4+ or CD8+ T_CM,_ T_EM_, and naïve T cells).

#### Pathway analyses

Further DEG lists characterizing the detectable and undetectable reservoir groups within cell subsets from RV254 were used to perform a multiple gene list analyses in Metascape to acquire the top 20 representative terms of the most significant enriched pathways ^63^. The genes comprising each of these 20 pathways were used as input lists to perform Gene Set Enrichment Analysis (GSEA) ^64^ when comparing the detectable and undetectable groups, along with an average expression matrix of all genes within each cell subset for each participant that was generated from the single-cell data. The GSEA results were filtered by normalized enrichment score (NES) ≥|1.4|, P < 0.001. For WGCNA-based pathway analyses, the CD14+ monocyte cell subset of the RV254 cohort Seurat object was used as input for coexpression analyses implemented in the single-cell R package, hdWGCNA ^65,66^. Metacells and a signed network were constructed within participants using non-default parameters (k=25, max_shared=10 and soft power=9). The top 25 hub genes for each of the resulting modules were used as a feature set for Seurat’s AddModuleScore to generate a score for each module within each cell. The Mann-Whitney U test was used to compare expression of the module scores between cells in the detectable and undetectable reservoir categories. This module scoring and testing method for the same sets of genes from RV254 was applied to the CD14+ monocyte cell subset in the independent A5354 cohort. Average scaled expression of the 25 hub genes from the M3 module containing *IL1B* within both cohorts was used as input for the ComplexHeatmap tool ^67^. Similarly, gene modules were identified in total CD4+ memory T cells and gene ontology analyses were performed using Enrichr ^68^. The 25 hub genes for the M3 CD14+ monocyte module were used as input in a protein STRING DB pathway analysis ^69^. The disconnected nodes were removed and the resulting network was investigated for degree of connectedness and visualized in Cytoscape ^70^.

### Statistical analyses

The associations between 117 phenotypic flow cytometry population frequencies and reservoir size were assessed by univariate linear regression models and corrected for multiple testing using false discovery rate (FDR). Exploratory analyses including multiple regressions without adjusting for significance were also performed to evaluate the relationship between the reservoir size as the response variable and two explanatory variables: *THBS1*/*IL1B* and each flow cytometry cell population. Finally, multiple regression models were fitted with two-way interaction terms between *THBS1*/*IL1B* and each phenotypic population marker, to test whether the effect of *THBS1*/*IL1B* on decreased reservoir size differed depending on the frequencies of individual cell subsets. Interaction plots for *THBS1*/*IL1B* were made to illustrate how the relationship between *THBS1*/*IL1B* expression and reservoir size changes with different frequencies of combined CD4+ T_CM_. The overall fitness of the simple regression models of the combined CD4+ T_CM_ population was evaluated using the coefficient of determination, R-squared value (R^2^), and Root Mean Squared Error (RMSE). For multiple linear regression of the CD4+ T_CM_ cells, the goodness-of-fit was measured using both R^2^ and Adjusted-R^2^ along with RMSE. The prediction error of the combined CD4+ T_CM_ cell models was estimated using Leave One Out Cross Validation (LOOCV) and the test RMSE value was reported. Assessment of model diagnostics (Q-Q plot for normality, residuals vs. fitted values for homoscedasticity, leverage plots for influential observations, variance inflation factors for multicollinearity; not shown) showed that the assumptions of the linear models were reasonable after removing one outlier. All explanatory variables for all regression analyses were mean centered.

All paired comparisons were performed by paired T test, when the assumptions were met, or the Wilcoxon signed-rank test, while unpaired comparisons were performed by the Mann-Whitney U test. Correlations were performed by Spearman’s rank correlation coefficient. A two-sided P value of <0.05 was considered statistically significant for all statistical analyses. Bonferroni or FDR corrections were applied for multiple testing when appropriate. All descriptive and inferential statistical analyses were performed using R 3.4.1 GUI 1.70 build (7375) v3.0 and higher, and GraphPad Prism 8.0 statistical software packages (GraphPad Software, La Jolla CA).

## Acknowledgements

We would like to thank Dr. Nicolas Chomont, University of Montreal, for supporting efforts to setup the HIV DNA assay at MHRP. Lilia Mei Bose and Hasset Tibebe, American University assisted with functional data analyses. We acknowledge Joseph Puleo, Summer Zheng and Justin Ritz, CBAR, Boston for providing IPDA data. The MR1 tetramer technology was developed jointly by Dr. James McCluskey, Dr. Jamie Rossjohn, and Dr. David Fairlie, and the material was produced by the NIH Tetramer Core Facility as permitted to be distributed by the University of Melbourne. We thank the participants and staff of the RV254/SEARCH010 and ACTG A5354 cohorts. The views expressed are those of the authors and should not be construed to represent the positions or views of the U.S. Army or the U.S. Department of Defense (DOD), the U.S. Centers for Disease Control and Prevention, the U.S. Public Health Service, or the U.S. Government.

## Data availability

All scRNA-seq gene expression data has been submitted to the GEO repository with accession number (GSE220790, GSE256089). Code is available in the Figshare database.

## List of Supplementary Materials

Figures S1 to S11

Tables S1 to S5

Data files 1-2

## SUPPLEMENTARY DATA (LEGENDS)

**Figure S1. Expression of select lineage markers in specific cell subsets**. Expression of known lineage marker genes is localized to certain regions on the scRNA-seq UMAP and enables cell type assignment of clusters.

**Figure S2. Clustering and frequency comparisons of scRNA-seq immune cell subsets.** scRNA-seq shows similar A) spatial patterns of cell populations and B) frequencies of cells between detectable (red) and undetectable (teal) reservoir groups. Significance was determined by the Mann-Whitney U test. * nominal p < 0.05

**Figure S3. T cell receptor and B cell receptor clonal diversity and isotype distribution.** Clonal diversity of **A)** indicated T cell populations and **B)** memory and naive B cell populations. Significance was determined by the Mann-Whitney U test. **C)** BCR isotype distribution of each donor within naïve and memory B cells.

**Figure S4. Gene expression in CD14+ monocytes across participants in RV254.** Gene expression in CD14+ monocytes of *IL1B* and *THBS1* genes across participants based on categorization of total HIV DNA measured from A) total PBMC and B) CD4+ T cells. Black circles represent the median values, and vertical lines indicate the interquartile range. Teal: undetectable reservoir; red: detectable reservoir. Three datapoints were missing due to technical differences and insufficient sample to perform the HIV DNA assay in sorted CD4+ T cells. C, D) Total HIV DNA associations when examining change in reservoir decay from week 0 (AHI) to 48 weeks after ART initiation with C) number of significant DEG and D) top genes associating with reservoir decay. E, F) Correlation of total HIV DNA with E) *THBS1* and F) *IL1B* gene expression in CD14+ monocytes.

**Figure S5. No differences in frequencies of CD4+ T_CM_ cell subsets between reservoir groups**. A) Gating strategy for the identification of CD4+ T_CM_ cell subsets from the RV254 participants. B) DR-CD4+ T_CM_, C) PD-1-CD4+ T_CM_ and D) CD4+ T_CM_. Significance was determined by the Mann-Whitney U test. Teal: undetectable reservoir; red: detectable reservoir.

**Figure S6. Pathway analyses identify distinct signatures associated with reservoir size.** A) WGCNA dendrogram shows modules of coexpressed genes in memory CD4+ T cells from the RV254 Thai study. B) The M4 and M9 WGCNA modules were enriched in cells from participants with undetectable reservoir (N=8) compared to detectable reservoir (N=6) based on the top 25 hub genes in the module. C) Using the same module hub genes found in RV254, these modules were enriched in cells from the undetectable reservoir participants in the A5354 cohort when HIV DNA levels were grouped categorically (detectable=12, undetectable=11). Enrichment analysis of D) M4 and E) M9 hub genes shows pathways significantly enriched (FDR < 0.05) in the two modules and the genes contributing to enrichment in their respective modules.

**Figure S7. Effects of HIV infection on in vitro CD4+ T cell memory phenotypes**. A) Gating strategy. B) Bar plot depicts the mean percentages of CD45RO-CCR7-Naive (Naive), CD45RO-CCR7+ Effector (Eff), and CD45RO+ memory (Mem) CD4+ T cell populations in total CD4+ T cells infected or not with HIV, as determined by intracellular p24 staining. C) Pie chart illustrates the mean percentages of memory subsets T_CM_, T_TM_, and T_EM_ within total CD45RO+ memory CD4+ T cells. * p<0.05.

**Figure S8. Effects of recombinant IL1B on HIV replication in PBMC cultures.** Time course plots of the infectious virus and p24 antigen yields of NL4-3 or CH058 in 5 individual donors. PBMC were isolated from buffy coats and incubated for 3 days, with PHA and IL-2, when they were infected with NL4-3 or CH058. IL1B (10 ng/ml) was added 2 days prior to infection (2d-), at the same time (0d), or 2 days post-infection (2d+). Supernatants were collected at designated time points and assessed for infectious virus and p24 antigen.

**Figure S9: Effect of IL1B on cell viability**. Isolated PBMCs were treated and infected as in Figures 5D (A) and S8 (B), respectively. On day 3 post-infection cells were harvested, stained with fixable viability dye and FACs analyzed for cell viability. Each dot represents a single donor.

**Figure S10. Effects of recombinant IL1B on in vitro CD4+ T cell memory phenotypes.** A) Gating strategy. B) Mean percentages of T_CM_, T_TM_, and T_EM_ subsets within total memory CD4+ T cells were determined following dose-dependent IL1B treatment. Statistical significance was assessed using the Mann-Whitney U test, based on comparing each treated or infected sample to its respective untreated or uninfected control. ** p<0.01.

**Figure S11. Working model of potential mechanisms for IL1B effects on reservoir size**. We hypothesize that IL1B may affect the latent HIV reservoir by 1) acting as a natural LRA, 2) contributing to reduced seeding of the reservoir, and 3) changing the composition of CD4+ T cell subsets.

**Supplementary table S1**. Different cell populations as detected in scRNA-seq analyses

**Supplementary table S2**. Differentially expressed genes in each cell subset (log fold change >|0.25|, Bonferroni p<0.05).

**Supplementary table S3**. Top 20 enriched pathways and processes.

**Supplementary table S4**. Enriched pathways for each reservoir phenotype stratified by cell subset (NES ≥|1.4|, p<0.001)

**Supplementary table S5**. Associations of increased *THBS1*/*IL1B* expression with cell subset population frequencies obtained by flow cytometry, and their effects on reservoir size

**Extended table 1**. Demographics and clinical data of participants from the study

**Extended table 2**. List of antibodies.

